# The Potassium Channel Subunit K_v_β1 is Required for Synaptic Facilitation

**DOI:** 10.1101/2020.02.05.935163

**Authors:** In Ha Cho, Lauren C. Panzera, Morven Chin, Scott A. Alpizar, Michael B. Hoppa

## Abstract

Analysis of the presynaptic action potential’s (AP_syn_) role in synaptic facilitation in hippocampal pyramidal neurons has been difficult due to size limitations of axons. We overcame these size barriers by combining high resolution optical recordings of membrane potential, exocytosis and Ca^2+^ in cultured hippocampal neurons. These recordings revealed a critical and selective role for K_v_1 channel inactivation in synaptic facilitation of excitatory hippocampal neurons. Presynaptic K_v_1 channel inactivation was mediated by the K_v_β1 subunit, and had a surprisingly rapid onset that was readily apparent even in brief physiological stimulation paradigms including paired-pulse stimulation. Genetic depletion of K_v_β1 blocked all broadening of the AP_syn_ during high frequency stimulation and eliminated synaptic facilitation without altering the initial probability of vesicle release. Thus using all quantitative optical measurements of presynaptic physiology, we reveal a critical role for presynaptic K_v_ channels in synaptic facilitation at small presynaptic terminals of the hippocampal neurons upstream of exocytic machinery.

**Significance:** Nerve terminals generally engage in two opposite and essential forms of synaptic plasticity (facilitation or depression) during high frequency stimulation that play critical roles in learning and memory. Measurements of the electrical impulses (action potentials) underlying these two forms of plasticity has been difficult in small nerve terminals due to their size. In this study we deployed a combination of optical measurements of vesicle fusion and membrane voltage to overcome this previous size barrier. Here, we found a unique molecular composition of Kv1 channel β-subunits that causes broadening of the presynaptic action essential to synaptic facilitation. Disruption of the K_v_β1 inactivation mechanism switches excitatory nerve terminals into a depressive state, without any disruption to initial probability of vesicle fusion.

## Introduction

The action potential (AP) firing pattern or “spike code” as typically measured from the soma is a gold standard for neural excitability within circuits. However, at presynaptic terminals the quantitative relationship between the input AP spike code and the magnitude of exocytosis, or vesicle fusion events per AP, can rapidly change due to stimulation frequency or firing pattern. Increased firing frequency can significantly increase the number of vesicles that fuse from an identical number of APs. This phenomenon is known as short-term synaptic facilitation, which can significantly enhance information transfer at synapses influencing several aspects of learning and memory (1). Thus, it is important to completely understand the underlying molecular and cellular mechanisms of synaptic facilitation.

A critical initial step in exocytosis is the arrival of AP_syn_at boutons, whose waveform can exhibit plasticity based on firing frequency. Repetitive firing may cause inactivation of axonal voltage-gated sodium channels (Na_v_) and voltage-gated potassium (K_v_) channels that control the depolarization and hyperpolarization of the waveform respectively. K_v_ inactivation primarily leads to an increase in AP width or broadening (2–9). The width of the AP_syn_ controls the fraction of time that Ca^2+^ channels open and the driving force of Ca^2+^ entry (10). The highly nonlinear influence of Ca^2+^ on exocytosis (11, 12) thus dictates that modest AP_syn_ broadening has the potential to critically impact synaptic facilitation (13–15). Indeed, AP_syn_ broadening during repetitive firing has been demonstrated to cause the facilitation of exocytosis in the pituitary nerve (3), dorsal root ganglion (16) and mossy fiber bouton (2), all due to K_v_ channel inactivation. However, the AP_syn_ waveform in Purkinje Cells has also been shown to undergo frequency-dependent decreases in amplitude that substantially contributes to synaptic depression (17). Therefore, it is best to consider the AP_syn_ as a plastic signal that can powerfully modulate exocytosis bidirectionally, rather than a digital spike soley acting as an initiation signal. We therefore reason that the somatic AP has proven to be a poor predictor of exocytosis magnitude as a result of a failure to resolve the AP_syn_ waveform and its molecular regulators in the majority of brain regions.

Investigating the molecular regulation of AP_syn_ in the common *en passant* nerve terminals of the cortex and hippocampus remains elusive due to the small size of these structures (<1 μm), which makes them inaccessible for whole-cell patch clamp recording. Our group has overcome this barrier by pioneering the use of genetically-encoded rhodopsin-based voltage indicators to measure AP_syn_ in *en passant* terminals of inhibitory and excitatory hippocampal neurons. Here we demonstrate a striking contrast between faciltating excitatory and depressing inhibitory nerve terminals in the hippocampus. We discovered that excitatory terminals are uniquely enriched with a comination of K_v_1.1/1.2 heteromers and K_v_β1 subunits. This combination of K_v_ subunits causes rapid AP_syn_ broadening during brief periods of high-frequency firing. This broadening was essential for enabling synaptic facilitation without altering initial exocytosis. Taken together, these results suggest that the molecular control of presynaptic K_v_ channel inactivation is an important modulator of synaptic facilitation upstream of vesicle release machinery.

## Results

Previously, our measurements of the AP_syn_ in hippocampal neurons found a very high ratio of K_v_ to Na_v_ channels, with K_v_1.1/1.2 channels dominating repolarization (18). We measured the sensitivity of exocytosis to changes in K_v_1.1/1.2 conductance using an optical probe of exocytosis (vGLUT1-pHluorin; vG-pH)(12, 19). Blockade of K_v_1.1/1.2 channels by application of dendrotoxin-κ (DTX) greatly enhanced exocytosis in excitatory hippocampal nerve terminals by 61±18% when stimulated with 1 AP **(Figure 1A-B)**. The sensitivity of exocytosis to DTX application was mirrored in the optical recordings of the AP_syn_ from neurons expressing the indicator QuasAr (20), with a characteristic broadening as measured by the full-width at half max (FWHM) of the waveform (30±4%) **(Figure 1C-D)** in agreement with previous findings (18). We measured this phenomenon in both excitatory and inhibitory neurons of the hippocampus without prior knowledge of their identity. Interestingly, the inhibitory neurons did not display any sensitivity to DTX-treatment as assayed by exocytosis (**Figure 1E-F**). Furthermore, the AP_syn_ waveform displayed no changes in amplitude or width from DTX treatment (**Figure 1G-H**). We confirmed the identity of neurons after measuring exocytosis or AP_syn_ waveform as excitatory or inhibitory using a fluorescent antibody directed against the lumenal domain of the vesicular GABA transporter (vGAT) as previously described (21) with example recordings for both cell types shown in **Figures 1I-J**. Taken together these experiments demonstrate a very selective enrichment of presynaptic enrichment of K_v_1 channels in excitatory nerve terminals of hippocampal neurons.

**Figure 1.**
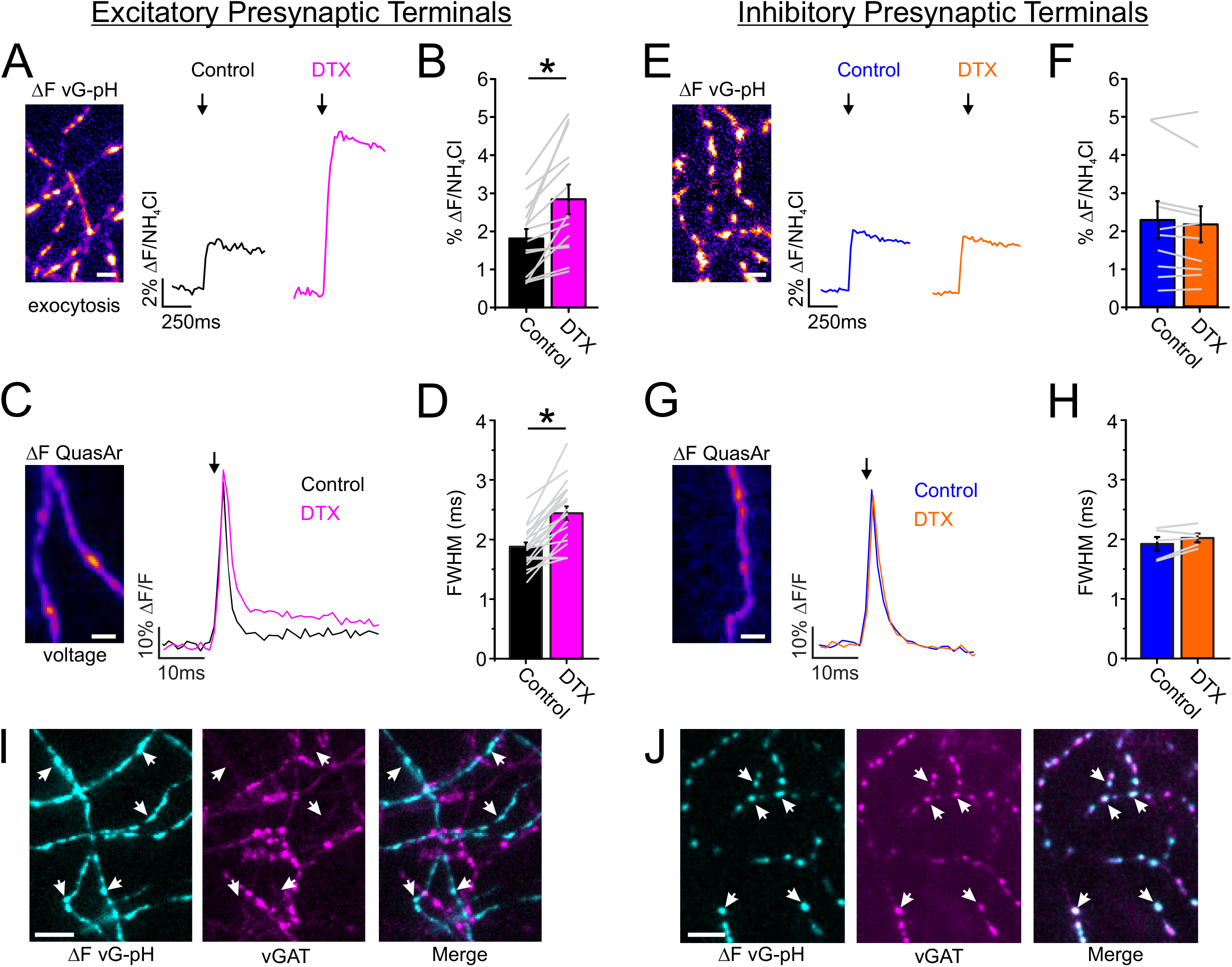
K_v_1 channels were expressed exclusively in excitatory presynaptic terminals. (**A-D**) Measurements of exocytosis using vG-pH (**A-B**) for both control (black) and DTX-treated (magenta) excitatory neurons; Arrow indicates when stimulation was applied; Example of recording of voltage using QuasAr (**C**) and corresponding FWHM (**D**);. Scale bar, 10 μm. (vG-pH, n = 16 cells, *p < 0.001, Paired t-test; QuasAr, n = 20 cells, *p < 0.01, Paired t-test). (**E-H**) Measurements of exocytosis using vG-pH (**E-F**) in control (blue) and DTX-treated (orange) inhibitory neurons. Representative recording of voltage using Quasar (**G**) and the corresponding averaged FWHM (**H**). Arrows indicate when stimulation was applied. (vG-pH, n = 10 cells; QuasAr, n = 5 cells). (**I-J**) Representative images of vGAT antibody live staining in excitatory terminals (**I**) and inhibitory terminals (**J**). Co-localization of vGAT staining signal with active synapses marked by vG-pH response indicated by the white arrows. Scale bar, 10 μm.

We hypothesized that terminals enriched with K_v_1.1/1.2 channels might exhibit AP_syn_ broadening during high frequency (>10 Hz) stimulation due to K_v_1 channel inactivation. We observed a robust (39±4%) broadening during a train of stimulation with 100 APs stimulated at 50 Hz (**Figure 2A**), with example binned recordings of the AP_syn_ shown in **Figure 2B** and the corresponding quantifications of the FWHM in **Figure 2C**. Next, we examined if AP_syn_ broadening also took place in short paired-pulse stimulation protocols associated with synaptic facilitation. We measured the FWHM of the AP_syn_ waveform during paired-pulse stimulation to compare how stable the shape of the waveform is across basal firing rates (4-10 Hz) and those typically associated with facilitation (50 Hz) (22, 23) (**Figure 2D-E**). The FWHM paired-pulse ratio is plotted as a function of the inter-spike interval in **Figure 2F** demonstrating that AP_syn_ broadening is reliably triggered by stimulation frequencies of 50 Hz. These results suggest that terminals enriched with K_v_1.1/1.2 channels in presynaptic terminals undergo frequency-dependent broadening of the AP_syn_ even in minimal conditions of paired-pulse stimulation that could influence vesicle fusion.

**Figure 2.**
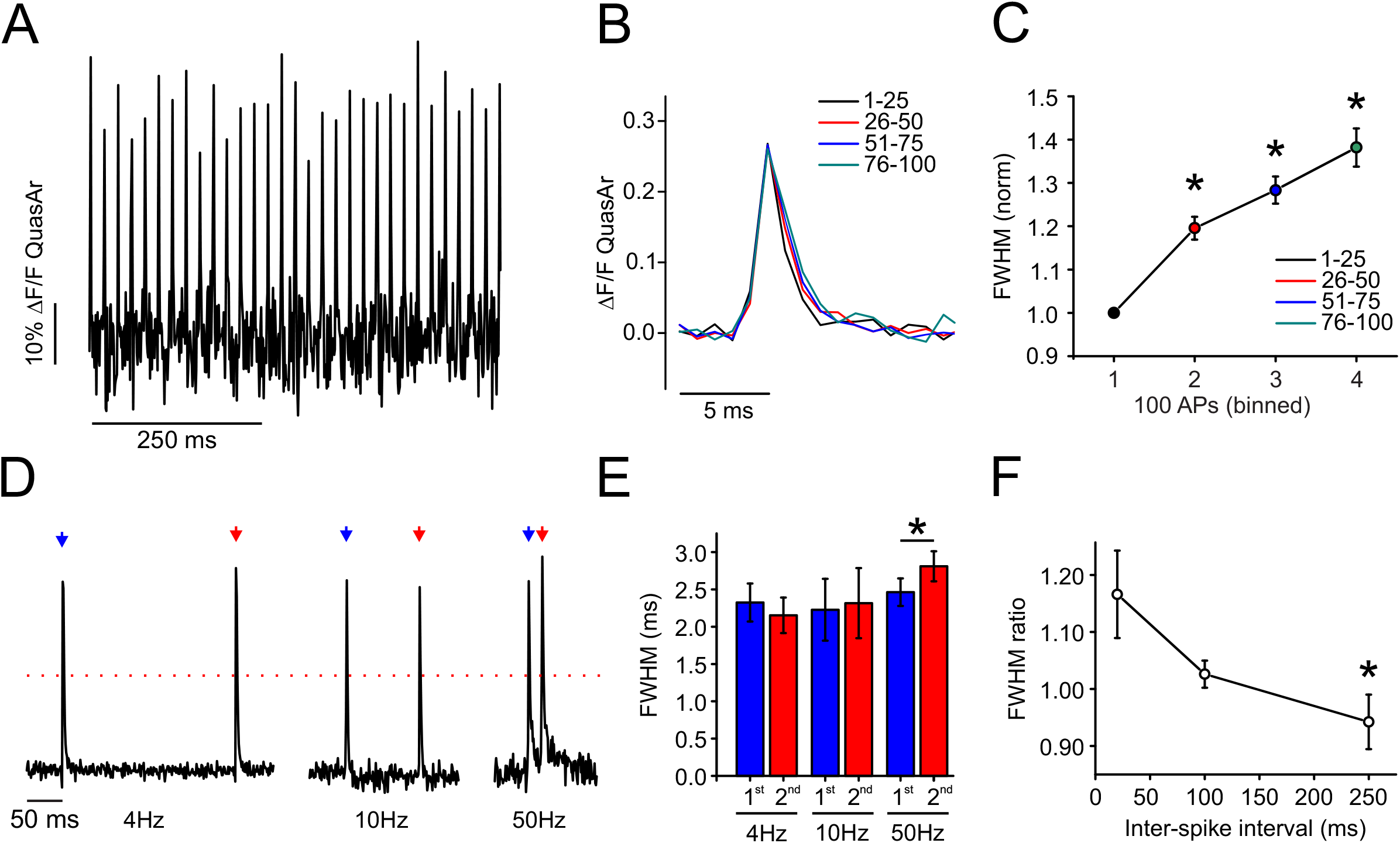
Activity-dependent broadening of AP_syn_ in excitatory nerve terminals of hippocampal neurons. (**A**) Example AP_syn_ traces evoked by 50 Hz stimulation. (**B**) Representative average AP traces from four separate bins of 25 APs during a 100 AP stimulation train delivered at 50 Hz. (**C**) Plot of normalized FWHM for four separate bins from excitatory cells. Black asterisk indicates significance relative to 1^st^ AP bin in excitatory cells (n = 39 cells, *p < 0.05, ANOVA with Tukey’s post hoc comparisons). (**D**) Representative measurement of QuasAr with 2 AP at 4 Hz, 10 Hz or 50 Hz stimulation in excitatory neurons. Peaks are normalized to the first peak of each measurment. Blue arrow indicates the first stimulation and red arrow indicates the second stimulation. Red dashed line represents the half maximum of the peak where the width was measured. (**E**) Average FWHM for the first (blue) and second (red) AP waveform in paired-pulse stimulation of excitatory neurons (n = 10 cells for 4 Hz condition; n = 8 cells for 10 Hz condition; n = 13 cells for 50 Hz; *p < 0.05, Paired t-test). Error bars indicate mean ± SEM. (**F**) Average FWHM ratio (2^nd^/1^st^ AP) is plotted as a function of a stimulation interval (*p < 0.05, ANOVA with Tukey’s post hoc comparisons).

While many changes in ionic conductances could underly the rapid broadening of AP_syn_ during stimulation, the most suggestive possibility from the previous experiments was that frequency-dependent K_v_1.1/1.2 channel inactivation was responsible for broadening. The dominant mechanism of K_v_1 family channel inactivation is the “ball-and-chain” mechanism, in which the N-terminal structures of either the K^+^ channel’s α or β subunits occlude the channel pore from the cytosol (24–26) (**Figure 3A**). K_v_1.1/1.2 channels are known to most prominently undergo inactivation when associated with cytosolic β subunits (27). As such, we investigated the role of the K_v_β1 subunit for AP broadening using shRNA (**Figure 3B-C**). We combined shRNA targeting the K_v_β1 subunits with QuasAr to determine the involvement of K_v_1.1/1.2 inactivation through the β-subunit in the broadening of the AP_syn_ during paired-pulse stimulation. The average waveforms are shown for excitatory neurons expressing scrambled shRNA (**WT; Figure 3D**) or shRNA directed against K_v_β1 (**K_v_β1KD; Figure 3E**). We also combined K_v_β1KD with expression of a human variant of K_v_β1 to *rescue* KD expression levels and check for off-target effects of KD (**hK_v_#x03B2;1; Figure 3F**). Genetic depletion of K_v_β1 not only stopped AP broadening, but also caused a small amount of narrowing (−7.0±2.6% in the FWHM) compared to control and rescue terminals (+10.4±4.1% and =8.9±2.6% in the FWHM) (**Figure 3G)**. This decrease in FWHM was accompanied by a more prominent hyperpolarization in KD neurons (**Figure 3D-F**) and an overall relative decrease (~11%) in presynaptic AP amplitude (**Figure 3H**), suggesting an enhancement of presynaptic K_v_1 currents without expression of K_v_β1 present. We used vG-pH to investigate the consequences of these changes in AP_syn_ broadening on the facilitation or depression of exocytosis (**Figure 3I**). We found that control (WT) terminals displayed a 42±10% increase in exocytosis (paired-pulse ratio of vG-pH is 1.42; facilitation > 1) when comparing stimulation from paired-pulses at 50 Hz to a single AP, but this enhancement or facilitation in vesicle fusion was completely abolished in K_v_β1 KD neurons (−9±8%; **Figure 3J**). We were curious if the inhibitory nerve terminals that do not contain K_v_1 channels underwent frequency dependent broadening of AP_syn_. We found that inhibitory nerve terminals mimicked the excitatory K_v_β1 KD terminals and exhibited paired-pulse narrowing of the AP_syn_ (**Figures S1A-B**). Furthermore, when we found that this APsyn narrowing was also accompanied by depression of neurotransmission during 50 Hz stimluation (vG-pH ratio < 1) akin to K_v_β1 KD excitatory neurons (**Figure S1C-D**). Taken together, these results indicate that K_v_β1 subunits play a critical role in AP broadening of excitatory nerve terminals during paired-pulse stimulation and suggest an important modulatory role for presynaptic K_v_1.1/1.2 currents in facilitating glutamate release.

**Figure 3.**
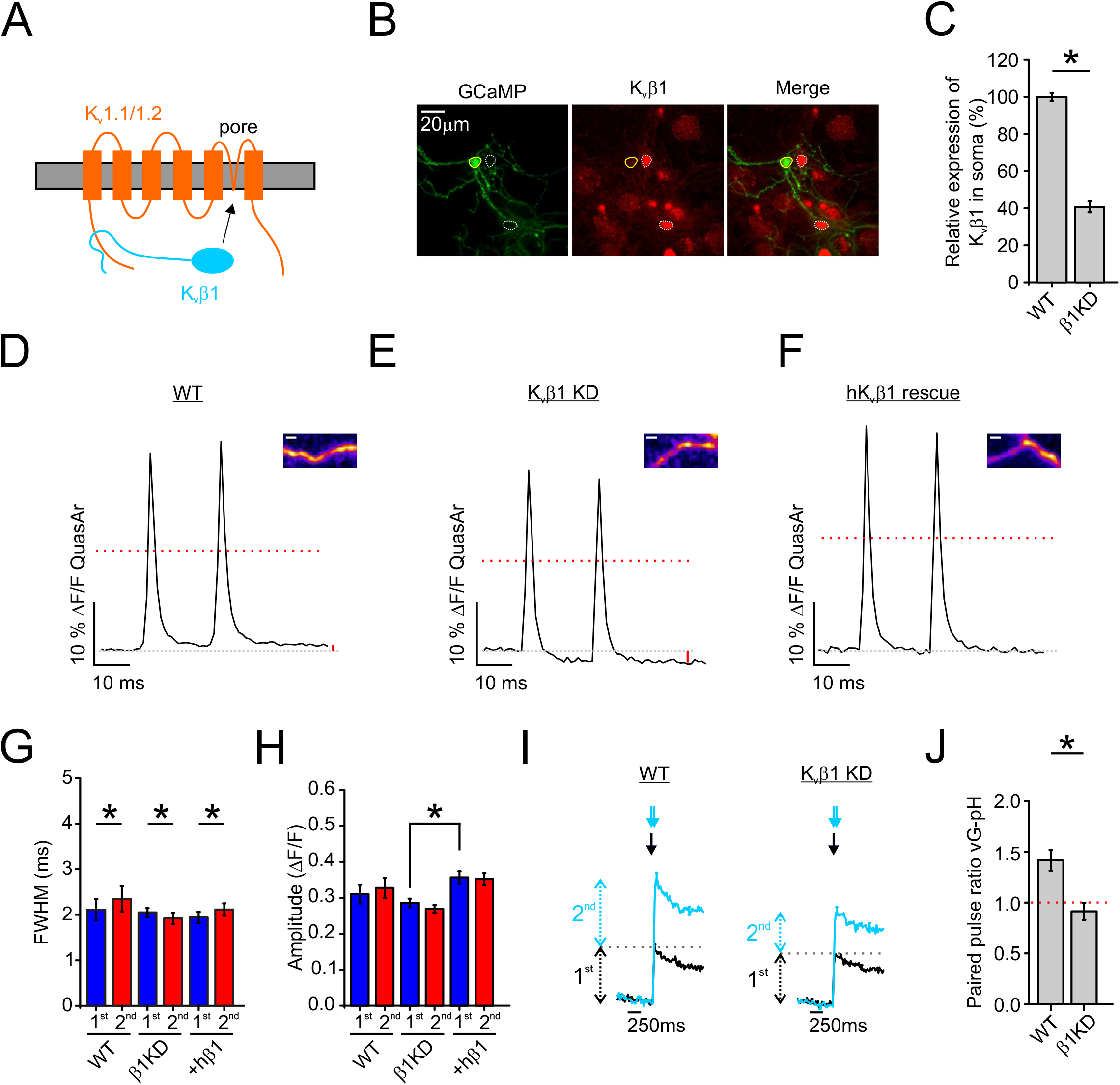
K_v_β1-induced K_v_1 inactivation is critical to AP_syn_ broadening with paired-pulse stimulation. **(A)** A cartoon of K_v_β1-induced inactivation of K_v_1.1/1.2 complexes. (**B**) Immunofluorescence staining for GCaMP (anti-GFP antibody) and K_v_β1 in primary cultured hippocampal neurons co-transfected with GCaMP and K_v_β1 shRNA. Solid lines indicate the soma of transfected neuron and dashed lines indicate that of untransfected neurons. Scale bar, 20 μm. (**C**) Quantification of relative expression of K_v_β1 in soma of K_v_β1 shRNA transfected neurons compared with control neurons (n = 50 for control; n = 19 cells for K_v_β1 shRNA transfected neurons; *p < 0.001, Student’s t test). (**D-F**) Average traces of AP waveforms in response to 2 AP at 50 Hz stimulation from control (**D**), K_v_β1 KD (**E**), and hK_v_β1 rescue (**F**) neurons. Insets provide a representative QuasAr ΔF image from each condition. Scale bar, 2 μm. (**G-H**) Average FWHM (**G**) and amplitude (**H**) for the first (blue) and second (red) AP waveform as shown in **D-F** (Control, n = 13 cells; K_v_β1 KD, n = 17 cells; hK_v_β1 rescue, n = 16 cells; *p < 0.05, Student’s t test for amplitude comparison between different conditions, Paired t-test for FWHM). (**I**) Average traces of exocytosis for 1 AP (black) or 2 AP delivered at 50 Hz (cyan) from control and K_v_β1 KD neurons as measured with vG-pH. (**J**) Average vG-pH ratio from control and K_v_β1 KD neurons (control, n = 8 cells; K_v_β1 KD, n = 9 cells; *p < 0.01, Student’s t-test). Error bars indicate mean ± SEM.

Hippocampal neurons typically fire in short bursts of APs during physiological conditions (28), so we next examined the contribution of K_v_β1-mediated K_v_1.1/1.2 inactivation during synaptic transmission consisting of ten electrical pulses delivered at 4 Hz or 50 Hz. We found that WT neurons displayed robust facilitation at 50 Hz stimulation (**Figure 4A)**. However, we observed no facilitation for K_v_β1 KD neurons, despite nearly identicle vesicle release at 4 Hz stimulation compared to WT neurons suggesting a selective impairment in facilitation (**Figure 4B-C**). Even under an extended protocol of fifty APs delivered at 50 Hz, no recovery is seen in exocytosis (**Figure S2**) in K_v_β1 KD neurons. We also measured the change of the FWHM of the AP waveform with minimal averaging (16 trials) for both control (WT; gray) and K_v_β1 KD neurons (KD; orange) for brief trains of 10 AP stimulation at 4 and 50 Hz (**Figure 4D-E**). Here again we found a complete loss of AP_syn_ broadening at 50 Hz stimulation for K_v_β1 KD neurons and a significant narrowing of the AP_syn_ 4 Hz stimulation (**Figure 4F**). We hypothesized that the loss of K_v_β1-mediated broadening of AP_syn_ could lead to the opening of fewer presynaptic Ca^2+^ channels and impaired net Ca^2+^ entry during high frequency stimulation. We tested this hypothesis using a cytosolic version of GCaMP6f and measured changes in presynaptic [Ca^2+^]_i_ during stimulation with 10 APs at 50 Hz for control (**gray; Figure 4G**) and K_v_β1 KD (**orange; Figure 4H**). Here we found that the presynaptic Ca^2+^ signal was strongly reduced (>50%) during trains of stimulation at 50 Hz for K_v_β1 KD neurons (**Figure 4I**). We also created bicistronic expression vectors to measure voltage paired with Ca^2+^ or vesicle fusion in single nerve terminals. These measurements had limited signal to noise under such restrictions, but showed a clear correlation between AP_syn_ broadening and vesicle fusion during paired-pulse stimulation, but not total Ca^2+^ in these limited measurements (**Figure S3A-L**). Critically, the magnitude of vesicle fusion elicited by a single AP in all conditions was not altered by the loss of K_v_β1 or stimulation at 4 Hz frquency suggesting that release probability and vesicle fusion machinery are intact, but that the AP_syn_ itself is a major modulator of faciltiation or depression. As such, these results indicate that minor changes in AP_syn_ broadening by loss of K_v_β1-mediated K_v_1 inactivation had a large cumulative impact on integrated [Ca^2+^]_i_ and synaptic facilitation during physiological patterns of activity.

**Figure 4.**
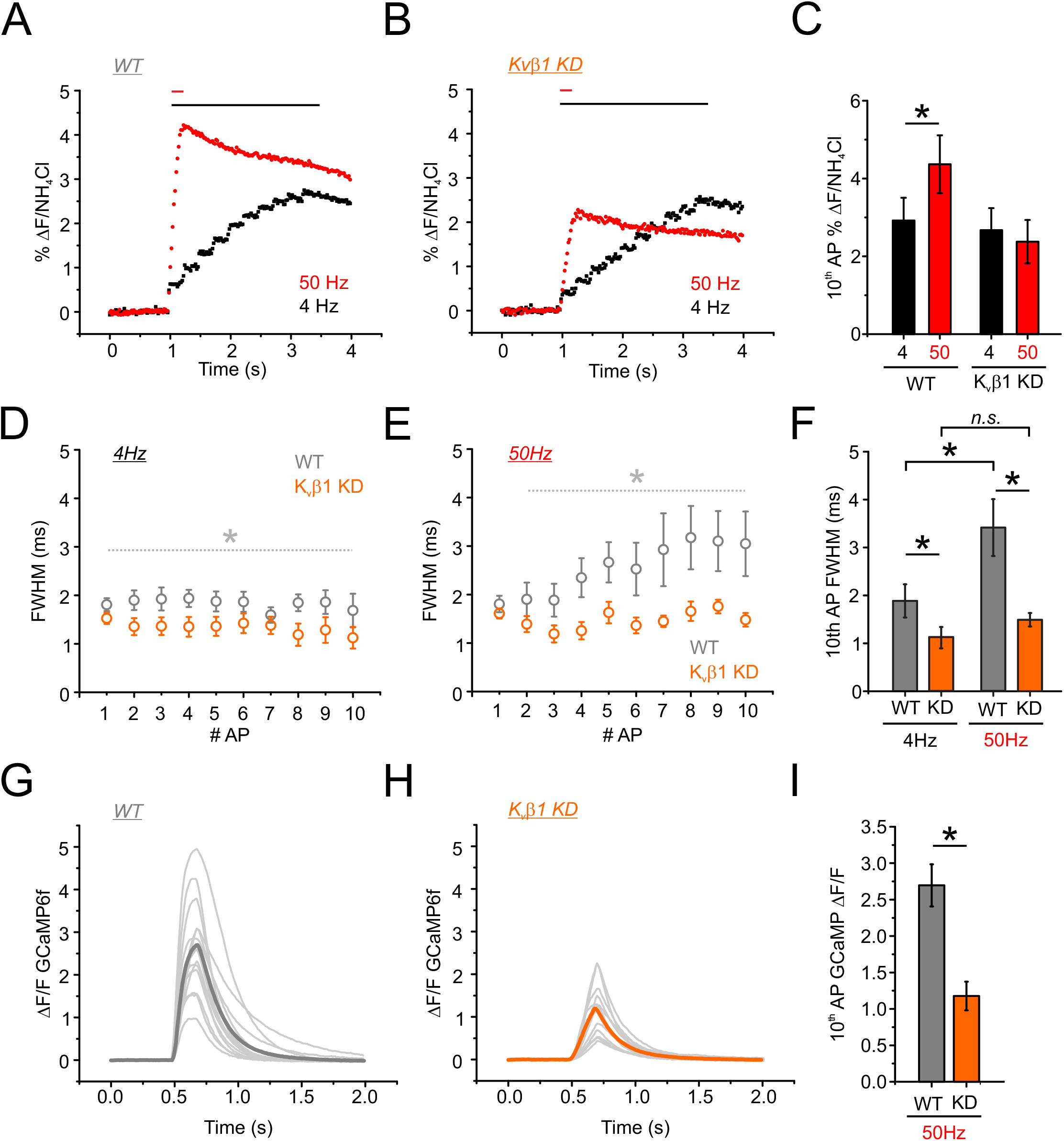
Synaptic facilitation is absent in K_v_β1 KD neurons. (**A-B**) Average traces of exocytosis for control (**A**) and K_v_β1 KD (**B**) in responses to 10 APs delivered at 4 Hz (black) or 50 Hz (red) as measured with vG-pH. Bars on top of the graphs indicate the duration of each stimulation. (**C**) Average fusion induced by the 10^th^ AP as a percentage of total vesicle fluorescences measured by application of NH_4_Cl. Neurons were stimulated with 10 AP at 4 Hz (black) or 50 Hz (red) (n = 11 cells,; K_v_β1 KD neurons, n = 8 cells; *p < 0.001, Paired-t test). (**D,E**) Optical recording of APs in neurons expressing QuasAr stimulated with 10 AP at 4 Hz (D) and 50 Hz (E). FWHM of each AP across the stimulus train from WT (gray) and K_v_β1 KD (orange) cells are displayed (n= 16 trials per cell; WT = 9 cells; K_v_β1 KD = 7 cells). (**F**) Quantification of the averaged FWHM of the 10^th^ AP from WT and K_v_β1 KD (n = 9 cells for WT; n = 7 cells for K_v_β1 KD; *p < 0.05, Student’s t test). Error bars indicate mean ± SEM. (**G-H**) Ca^2+^ influx was measured with GCaMP6f in control (G) and K_v_β1 KD (H) neurons. The light gray traces represent individual experiments with the averaged Ca^2+^ influx depicted in dark gray. (**I**) Quantification of the averaged GCaMP6f response of the 10^th^ AP from WT and K_v_β1 KD (n = 14 cells for WT; n = 11 cells for K_v_β1 KD; *p < 0.001, Student’s t test).

## Discussion

Our central finding is that a necessary mechanism of synaptic facilitation in excitatory hippocampal neurons is AP_syn_ broadening. We find the surprisingly rapid frequency-dependent broadening of AP_syn_ is enabled by a unique molecular combination of Kv1.1/1.2 channels with the K_v_β1 subunit. Indeed, this small broadening of the AP_syn_ mediated by K_v_β1 has a tremendous impact on synaptic transmission as the loss of the K_v_β1 subunit blocks synaptic facilitation without altering initial vesicle fusion (**Figure 3–4**). We believe that the conditions of AP_syn_ broadening works to facilitate exocytosis through a host of additional molecular interactions that minimally include downstream Ca^2+^ sensors and enzymes, but that K_v_1 inactivation represents a critical initial step to enable facilitation. This combination of Kv1 isoforms and subunits is not a ubiquitous system and even in cultured hippocampal neurons as inhibitory neurons even within the hippocampal culture demonstrated substantially different modulation of AP_syn_ and presynaptic K_v_ isoform enrichment (**Figure 2**). As such, we believe a detailed accounting of the channels responsible for the AP_syn_ shape across axons and associated terminals in various neural cell types could be an important contribution to our understanding of short-term synaptic plasticity and circuit dynamics. A central limitation of this study is that high-resolution optical measurement of AP_syn_ in hippocampal neurons are limited to *in vitro* measurements. Nevertheless, we believe this well-studied model for presynaptic function will translate into understanding how AP_syn_ relates to short-term plasticity and learning and memory *in vivo* such as those previously reported for the K_v_β1 knockout mouse (29).

Forms of synaptic enhancement, such as facilitation, augmentation, and post-tetanic potentiation, are usually attributed to effects of a residual elevation in presynaptic [Ca(^2+^)]_i_, acting on one or more molecular targets (Zucker 2002). In the latter regard, a critical specialized high-affinity Ca^2+^ sensor synaptotagmin 7 (Syt-7) (30) was identified to be required for facilitation in several regions of the brain including the hippocampus (28). However, subsequent studies have identified several other neural cell types that exhibit synaptic depression also have prominent levels of Syt-7 (31–34). Taken together, these experiments suggest Syt-7 produces facilitation in coordination with other molecular cascades that remain to be determined. Our data suggest that part of this coordination is upstream of Ca^2+^ sensing and vesicle release machinery. A widely accepted, but difficult-to-test model of facilitation attributes the enhancement of exocytosis to residual Ca^2+^ buildup during high frequency stimulation. *En passant* terminals like those in the cortex and hippocampus have been shown to have very efficient clearance of Ca^2+^ by diffusion thanks to abundant adjacent axonal volume unlike larger single terminals such as the Calyx of Held (35). One identified mechanism for this enhancment of intacellular Ca^2+^ during high-frequency stimulation is activity dependent activation of Ca_v_ channels (36). However this does not seem to be relevant in hippocampal neurons at physiological temperatures (37) like the ones we used in our experiments. We measured Ca^2+^ in control and K_v_β1 KD neurons and found that substantial build up of global [Ca^2+^]_i_ is strongly enhanced by AP_syn_ broadening when measured during physiological trains of stimulation (**Figure 4**). We speculate that this is especially critial in light of efficient Ca^2+^ extrusion coupled with the low overshoot (+7 mV) of the excitatory AP_syn_ that will typically only open a fraction of available presynaptic Ca^2+^ channels, a phenomenon we have previously identified in excitatory terminals of the hippocampus (18). Thus AP_syn_ broadening may be a particuarly efficient mechanism for facilitation in *en passant* boutons in general.

While we demonstrate that AP_syn_ broadening mediated by K_v_1 inactivation is essential for facilitation in excitatory hippocampal neurons, we do not believe it is conserved across all terminals as demonstrated by our measurements in inhibitory neurons (**Figure 2**). Interestingly, it appears that without K_v_β1 subunits, overall K_v_ currents are activating during high frequency stimulation, resulting in significant narrowing of AP_syn_ clearly observed for paired-pulse stimulation in inhibitory neurons as well as K_v_β1 KD excitatory neurons. Presynaptic K_v_ channel activation during high frequency stimulation was previously reported in the Calyx of Held (38) and may indeed be the default mode for many presynaptic K_v_ channels. This activation of K_v_ channels could be a very useful property for neurons that typically exhibit high firing rates such as PV neurons to depress exocytosis and to maintain a supply of vesicles as well as the timing of neurotransmitter release. This also seemed to be the case in Purkinje cell terminals that who frequency-dependent attenuation of AP_syn_ at larger terminals synapsing with deep cerebellar nuclei (17). The mechanisms that activate K_v_ channels are not fully resolved at present and may involve the recruitment of Ca^2+^ sensitive K^+^ channels such as SK and BK channels. What is clear is that K^+^ channel inactivation during physiological stimulation is not inherent and in the case of some K_v_1 channels (K_v_1.1 and 1.2 heteromers) requires binding partners. Here we identified the K_v_β1 subunit as a powerful modulator of exocytosis and synaptic facilitation. K_v_1 channel inactivation by the K_v_β subunit is well conserved with the respective fly homologues of *Shaker* and *Hyperkinetic* found in *Drosophila* to act in a similar manner (39). Additionally, impaired inactivation of K_v_1 channels was also identified as a presynaptic channelopathy of ataxia, which combined with our findings could suggest a broad importance across several circuits in the brain (40). Previous behavioral experiments in mouse identified the K_v_β1 subunits as critical for many memory tasks in K_v_β1 knockout mice (29), we believe our findings may be mechanistically helpful to understanding this phenotype. Slice recordings in this knockout mouse did not show impaired facilitation. However, we point out that a critical difference between these measurements and ours was the temperature of the recordings in slice were peformed at room temperature unlike what we report here at physiological temperatures (>34°C). We believe our data provide evidence that the AP_syn_ waveform is a critical modulator of synaptic facilitation in excitatory nerve terminals and that further study of presynaptic K^+^ channels is warranted across neuronal cell types.

## Acknowledgements

We would like to thank Andrew Coleman, Kelly Forest, and Amelia Ralowicz and for critical reading of the manuscript.

## Funding

This work was supported by the Esther A. and Joseph Klingenstein Fund (SAA, MBH), National Science Foundation’s Grant IOS 1750199 (IHC, MBH) and support from the Graduate Assistance in Areas of National Need GAANN Department of Education (SAA) P200A150059. NIH P20 NIGMS grant GM113132 (MBH).

## Materials and Methods

### Cell culture and transfection

Primary hippocampal neurons from postnatal day 1 Sprague Dawley rats of either sex were cultured as described previously (Cho 2017). Hippocampal CA1-CA3 regions were digested with trypsin for 5 min at room temperature and dissociated into single cells. Cells were seeded inside a 6 mm diameter cloning cylinder on polyornithine-coated coverslips. Plasmids were transfected into 5 - 6 DIV (Days in Vitro) neurons with calcium-phosphate precipitants. All animal protocols were approved by Dartmouth College’s Institutional Animal Care and Use Committee (IACUC protocol number 0002115).

### Plasmids

QuasAr constructs and vGlut-pHluorin were obtained as described previously (41), respectively. To measure membrane potential and Ca^2+^ influx/vesicle fusion in the same cell, we designed a QuasAr-P2A-GCaMP/QuasAr-P2A-vGlut-pHluorin by inserting QuasAr fused to P2A peptide synthesized using GeneArt Gene Synthesis (Invitrogen) and GCaMP/vGlut-pHluorin under the human Synapsin1 promoter, inducing independent expression of two different proteins in neurons. We validated that QuasAr and GCaMP/vGlut-pHluorin were co-expressed in the same cells through immunostaining assay and many repetitive experiments. To knock-down endogenous K_v_β1 expression, shRNA plasmids were obtained from OriGene against the following mRNA target sequences: GCTTGGTCATCACAACCAAACTCTACTGG. For rescue experiments in K_v_β1 knockdown neurons, human K_v_β1.1 fused to P2A peptide synthesized using GeneArt Gene Synthesis was inserted into pFCK-QuasAr plasmid to induce independent expression of QuasAr and human K_v_β1.1 in neurons.

### Antibodies and Reagents

Chicken polyclonal anti-GFP and rabbit polyclonal anti-mCherry were purchased from Invitrogen. Rabbit polyclonal anti-vGAT-Alexa 550 and mouse monoclonal anti-Synapsin1 were obtained from Synaptic Systems. Mouse monoclonal anti-K_v_β1.1 was purchased from Neuromab. Alexa Fluor 488-, 546-, and 647-conjugated goat anti-rabbit, anti-mouse, anti-Chicken and anti-guinea pig IgG were from Thermo Fisher Scientific. Dendrotoxin-k was purchased from Alomone (Israel).

### Live cell imaging

All live imaging experiments were set up as previously described (42, 43). Briefly, images were obtained using an Olympus microscope (IX-83) equipped with a 40x 1.35 NA oil immersion objective (UApoN40XO340-2) and captured with an IXON Ultra 897 EMCCD (Andor). Coverslips were mounted in a laminar-flow perfusion and stimulation chamber on the stage of the microscope. Cells were perfused continuously at a rate of 400 μl/min in Tyrode solution containing the following (in mM): 119 NaCl, 2.5 KCl, 2 CaCl2, 2 MgCl2, 25 HEPES, and 30 glucose with 10 μM CNQX and 50 μM AP5 during experiments. All experiments were performed at 34-35 °C with a custom-built objective heater.

For K_v_1 channel experiments, dendrodotoxin (DTX) was used at 100 nM concentrations. Cells were incubated with DTX in Tyrode solution for 1 min, followed by perfusion with normal Tyrode solution for additional 1 min and images were taken.

For measuring membrane potential, fluorescence of QuasAr was recorded with an exposure time of 980 μs and images were acquired at 1 kHz using an optomask (Cairn Research) to prevent light exposure of non-relevant pixels. Cells were illuminated by a 637 nm laser 70 - 120 mW (Coherent OBIS laser) with ZET635/20×, ET655lpm and ZT640rdc filters all obtained from Chroma. We repeated more than 100 trials to measure axonal AP waveform and averaged the signals, except 10 AP stimulation results in Figure 4 (16 trials). Timing of stimulation was delivered by counting frame numbers from a direct readout of the EMCCD rather than time itself for more exact synchronization using an arduino board and software custom manufactured by Sensorstar (Elkridge, MD).

For measuring Ca2+ influx and vesicle fusion, GCaMP6f and vGlut-pHluorin were collected with an exposure time of 20 ms and images were acquired at 50 Hz. Cells were illuminated by a 488 nm laser 6-8 mW (Coherent OBIS laser) with ET470/40×, ET525/50m, and T495lpxr filters (Chroma). We repeated more than 16 trials to measure single AP-induced responses and 6 trials to measure AP train stimulation-induced responses.

For differentiating inhibitory neurons from excitatory neurons, live vGAT antibody staining was performed using Alexa 550 pre-labeled vGAT antibody at the end of the experiments. Cells were stimulated by 1000 AP at 10 Hz with the antibody and followed by perfusion with normal Tyrode solution for 5 min. Fluorescence was captured using ET560/40×, ET630/75m, and T585lpxr filters (Chroma).

### Immunofluorescence

To confirm the co-expression of QuasAr and GCaMP or vGlut-pHluorin in the neurons, 14-17 DIV neurons were fixed with 4 % paraformaldehyde and 4% sucrose in PBS for 10 min, permeabilized with 0.2% Triton X-100 for 10 min and blocked by 5% goat serum in PBS for 30 min at room temperature. Cells were then incubated with the appropriate primary antibodies and visualized using Alexa Fluor-conjugated secondary antibodies in the PBS containing 5% goat serum. Images were obtained using a custom-made fluorescence microscope equipped with 40x oil-immersion objectives and filters described above.

### Image and data analysis

Images were analyzed in ImageJ and Fiji using custom-written plugins (http://rsb.info.nih.gov/ij/plugins/time-series.html). All statistical data are presented as means +/-SEM (n = number of neurons) and all experiments were performed on more than three independent cultures. For examining AP broadening with QuasAr measurement, we used Origin version 9.1. To obtain FWHM of each peak, half maximum of first peak was applied to every peak. For analyzing vesicle fusion of each condition, it was normalized to total number of vesicles measured by ammonium chloride treatment at the end of the experiments.

### Quantification and statistical analysis

Statistical analyses were performed in Excel and Origin. We used paired two sample for means t-test for paired results. Normally distributed data were processed with the Student’s t-test for two independent distributions and with a one-way ANOVA followed by Tukey’s post hoc comparison for comparing more than two groups to examine statistical significance. We specified the use of these tests and exact sample sizes in the figure legends for clarity.

## Supplementary Figure Legends

**Supplementary Figure 1.**
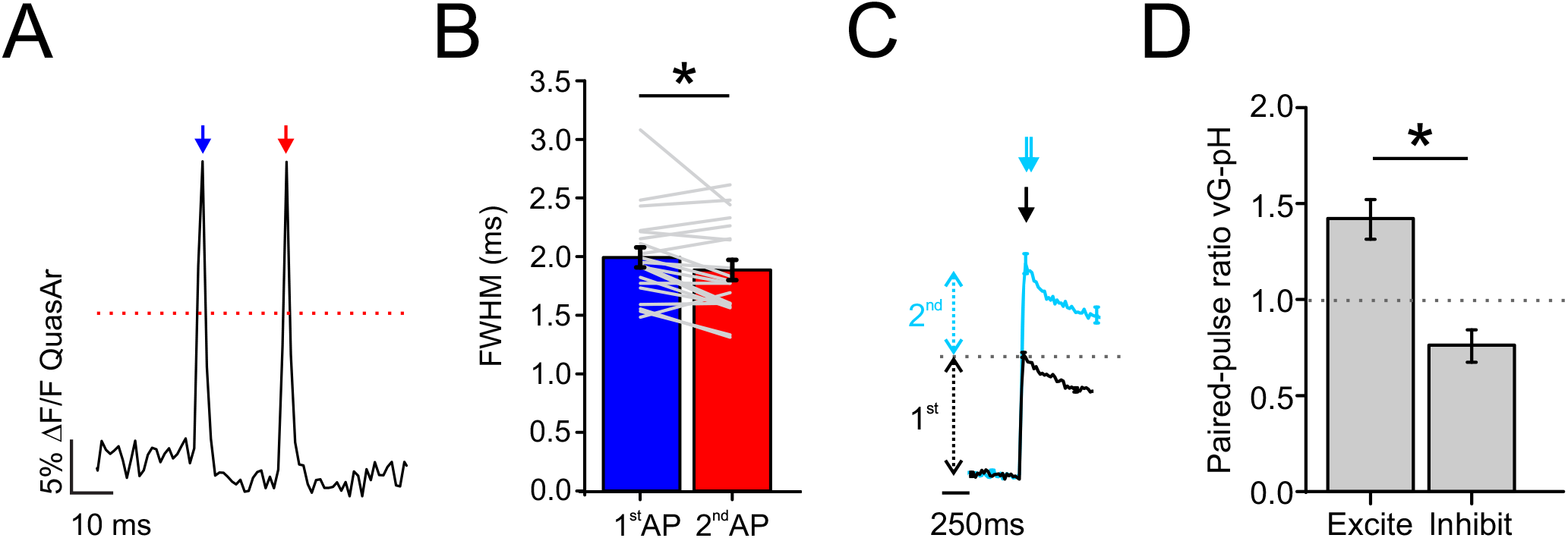
Presynaptic terminal of inhibitory neurons do not exhibit activity dependent broadening of AP_syn_ and synaptic facilitation by paired-pulse stimulation. **(A)** Representative measurement of voltage with QuasAr of 2 AP delivered at 50 Hz. Blue arrow indicates the first stimulation and red arrow indicates the second stimulation. Dashed red line represents the half maximum where the FWHM was measured. **(B)** Average FWHM for the first (blue) and second (red) AP waveform in paired-pulse stimulation at 50 Hz of inhibitory neurons (n = 20 individual cells, *p < 0.05, Paired t-test). **(C)** Average traces of vesicle fusion for 1 AP (black) or 2 AP delivered at 50 Hz (cyan) as measured with vG-pH. **(D)** Average vG-pH ratio from excitatory (reproduced from Figure 3J) and inhibitory neurons (n = 8 individual cells for inhibitory neurons, *p < 0.01, Student’s t-test).

**Supplementary Figure 2.**
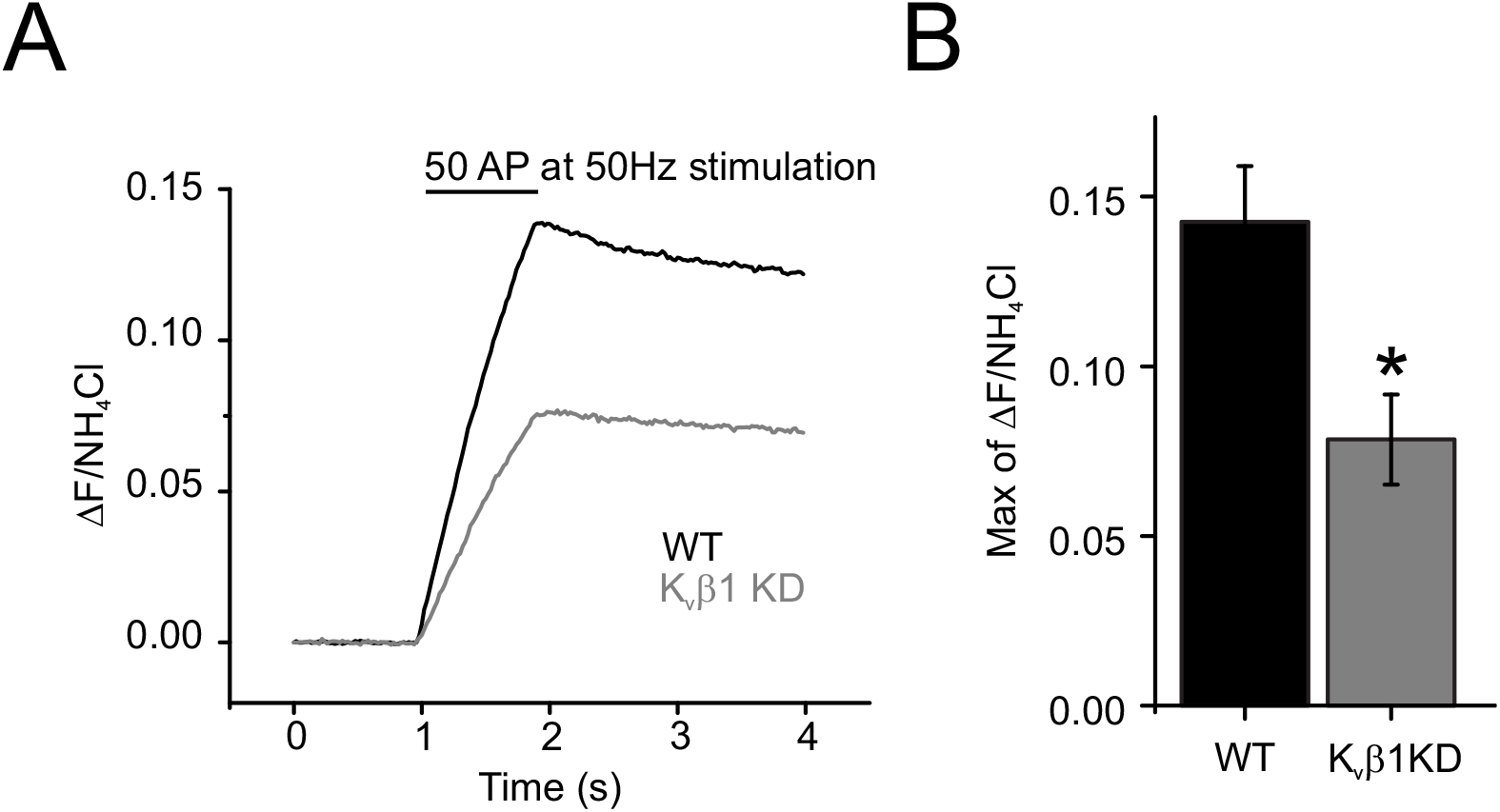
Impairment of synaptic facilitation during long trains of stimulation in K_v_β1 KD neurons. **(A)** Average traces of vesicle fusion in responses to a 50 AP at 50 Hz stimulation as measured with vG-pH from control (black) and K_v_β1 KD (gray). **(B)** Average max of % total vesicles (normalized to NH_4_Cl akalanization) for control and K_v_β1 KD neurons (n = 22 for control; n = 9 for K_v_β1 KD; *p < 0.01, Student-t test).

**Supplementary Figure 3.**
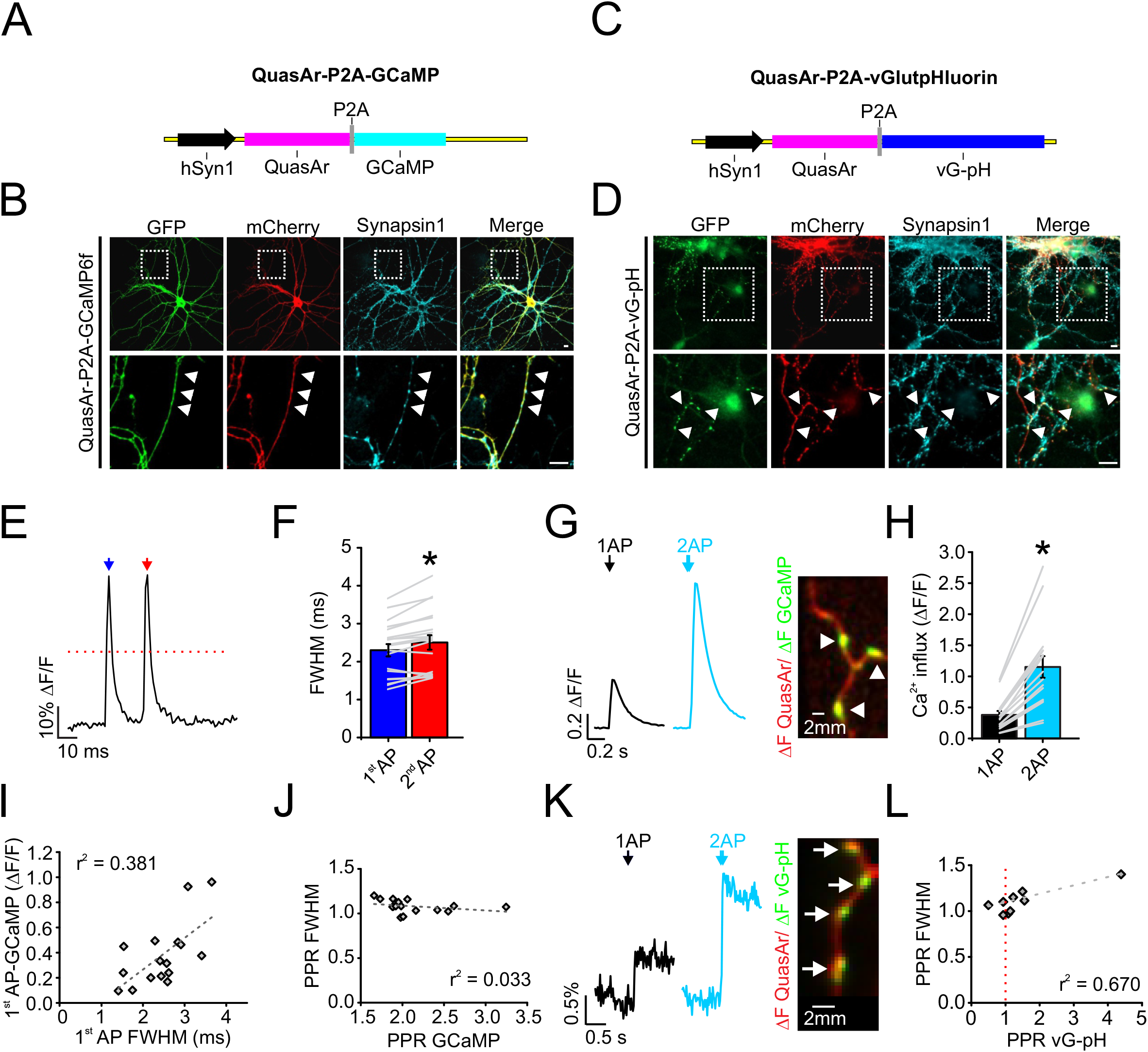
AP broadening by paired-pulse stimulation in excitatory neurons correlate with synaptic facilitation, but not with Ca^2+^ facilitation. **(A)** Schematic of vector describing bicistronic expression of QuasAr and GCaMP separated by a P2A peptide that cleaves the two proteins for separate localization. **(B)** Immunofluorescence of QuasAr-P2A-GCaMP expression in axons. GCaMP and QuasAr were stained with anti-GFP and anti-mCherry antibodies, respectively. QuasAr contains a mOrange-tag mutated to be non-fluorescent as to not have cross talk with the GFP signal. Synapsin1 staining was used for marking presynapses. Arrowheads mark presynaptic terminals. Scale bar, 10 μm. **(C)** Schematic of vector describing bicistronic expression of QuasAr and vG-pH separated by a P2A peptide that cleaves the two proteins for separate localization. **(D)** Immunofluorescence of QuasAr-P2A-vGpH expression in axons. vG-pH and QuasAr were stained with anti-GFP and anti-mCherry antibodies, respectively. **(E)** Representative measurement of QuasAr from axon with *en passant* synapses with 2 AP at 50 Hz stimulation in excitatory neurons. Blue arrow indicates the first stimulation and red arrow indicates the second stimulation. Red dashed line represents the half maximum where the width was measured. **(F)** Average FWHM for the first (blue) and second (red) AP waveform in paired-pulse stimulation of excitatory neurons (n = 19, *p < 0.001, Paired t-test). Error bars indicate mean ± SEM. **(G)** *Left,* Representative trace of GCaMP response to a single AP (blue) and paired-pulse stimulation (cyan). *Right,* Representative QuasAr ΔF (red) and GCaMP ΔF images (green) during the stimulation. Scale bar, 2 μm. **(H)** Average Ca^2+^ influx measured by GCaMP in paired-pulse stimulation (n = 16 individual cells, *p < 0.001, Paired t-test). **(I)** Correlation between 1^st^ AP induced Ca^2+^ influx and FWHM of 1^st^ AP. Linear fit is shown using a dashed line. **(J)** Correlation between paired-pulse ratio of GCaMP (PPR GCaMP) and FWHM (PPR FWHM) using a linear fit. **(K)** *Left,* Representative trace of vG-pH response to a single AP (black) and paired-pulse stimulation (cyan). *Right,* Representative QuasAr ΔF (red) and vG-pH ΔF images (green) during the stimulation. Scale bar, 2 μm. **(L)** Correlation using a linear fit between paired-pulse ratio of vG-pH (PPR vG-pH) and FWHM (PPR FWHM). Gray dashed line is the fitted line (n = 9). Vertical red dashed line indicates where vesicle fusion doubles in the second AP relative to the first AP.

